# Comparative analysis of primer-probe sets for the laboratory confirmation of SARS-CoV-2

**DOI:** 10.1101/2020.02.25.964775

**Authors:** Yu Jin Jung, Gun-Soo Park, Jun Hye Moon, Keunbon Ku, Seung-Hwa Beak, Seil Kim, Edmond Changkyun Park, Daeui Park, Jong-Hwan Lee, Cheol Woo Byeon, Joong Jin Lee, Jin-Soo Maeng, Seong Jun Kim, Seung Il Kim, Bum-Tae Kim, Min Jun Lee, Hong Gi Kim

## Abstract

Coronavirus disease 2019 (COVID-19) is newly emerging human infectious diseases, which is caused by Severe Acute Respiratory Syndrome Coronavirus 2 (SARS-CoV-2, also previously known as 2019-nCoV). Within two months of the outbreak, more than 80,000 cases of COVID-19 have been confirmed worldwide. Since the human to human transmission occurred easily and the human infection is rapidly increasing, the sensitive and early diagnosis is essential to prevent the global outbreak. Recently, World Health Organization (WHO) announced various primer and probe sets for SARS-CoV-2 previously developed in China, Germany, Hong Kong, Japan, Thailand, and USA. In this study, we compared the ability to detect SARS-CoV-2 RNA among the seven primer-probe sets for N gene and the three primer-probe sets for Orf1 gene. The result of the comparative analysis represented that the ‘2019-nCoV_N2, N3’ of USA and the ‘ORF1ab’ of China are the most sensitive primer-probe sets for N and Orf1 genes, respectively. Therefore, the appropriate combination from ORF1ab (China), 2019-nCoV_N2, N3 (USA), and NIID_2019-nCOV_N (Japan) sets should be selected for the sensitive and reliable laboratory confirmation of SARS-CoV-2.

## Introduction

Firstly informed to World Health Organization (WHO) on 31 December 2019, the current outbreak of Coronavirus Disease (COVID-19) involves 78,811 confirmed cases over 28 countries as of 23 February 2020 [1]. The majority of COVID-19 patients had pneumonia and showed symptoms include fever and cough [2, 3]. The genome sequence of causative novel coronavirus was shared through Global Initiative on Sharing All Influenza Data (GISAID) platform from 12 January 2020. The sequences of novel coronavirus (CoV) showed close similarity to that of severe acute respiratory syndrome-related coronaviruses (SARSr-CoV) and the virus uses ACE2 as the entry receptor like SARS-CoV [4-6]. The Coronavirus Study Group of the International Committee on Taxonomy of Viruses designated the virus as SARS-CoV-2 [7].

Molecular diagnosis of COVID-19 is currently carried out by one-step quantitative RT-PCR (qRT-PCR) targetting SARS-CoV-2 by which primers and probes being suggested by institutes of China, Germany, Hong Kong, Japan, Thailand, and USA were posted through WHO [8-10]. Clinical diagnosis methods including CT scan are also utilized to identify COVID-19 cases in Hubei province, China, from 13 February 2020 [11]. Although qRT-PCR assay served as a gold-standard method to detect respiratory infectious viruses such as SARS-CoV and MERS-CoV [12-15], current qRT-PCR assays targetting SARS-CoV-2 have some caveats. First, due to the high similarity of SARS-CoV-2 to SARS-CoV, primer-probe sets would cross-react. Second, the sensitivity of the assays may not enough to confirm suspicious patients in early time points after admission. Indeed, cases of positive CT scan results and negative RT-PCR results at initial presentation were reported [16]. The performance of molecular diagnosis might be dependent on primers, probes, and reagents. There have been no comparative results of the current qRT-PCR analysis for the molecular diagnosis of SARS-CoV-2.

In this present study, the qRT-PCR analysis was performed with previously reported primer-probe sets targeting RdRp/Orf1 and N region of SARS-CoV-2. This is the first comparative analysis of various primer-probe sets for the laboratory confirmation of SARS-CoV-2.

## Materials and methods

### Primer Information of qPCR

For the comparative analysis of laboratory confirmation for SARS-CoV-2, ten primer-probe sets were selected based on sequence information from the six different national institutions; the Centers for Disease Control and Prevention (CDC) (USA), Charité – Universitätsmedizin Berlin Institute of Virology (Germany), The University of Hong Kong (Hong Kong), National Institute of Infectious Disease, Department of virology III (Japan), China CDC (China), and National Institute of Health (Thailand). All of the DNA oligonucleotides were synthesized from Neoprobe (Daejeon, South Korea). The sequences of primer-probe sets and their locations at viral RNA (GenBank MN908947.3) were listed in Figure 1 and Table 1. Seven of the ten sets were derived from the N gene, and the other three sets were derived from Orf1 gene (RdRp, ORF 1b-Nsp14, and ORF 1-Nsp10). All DNA oligonucleotides were resuspended in nuclease-free water before use.

**Table 1.**
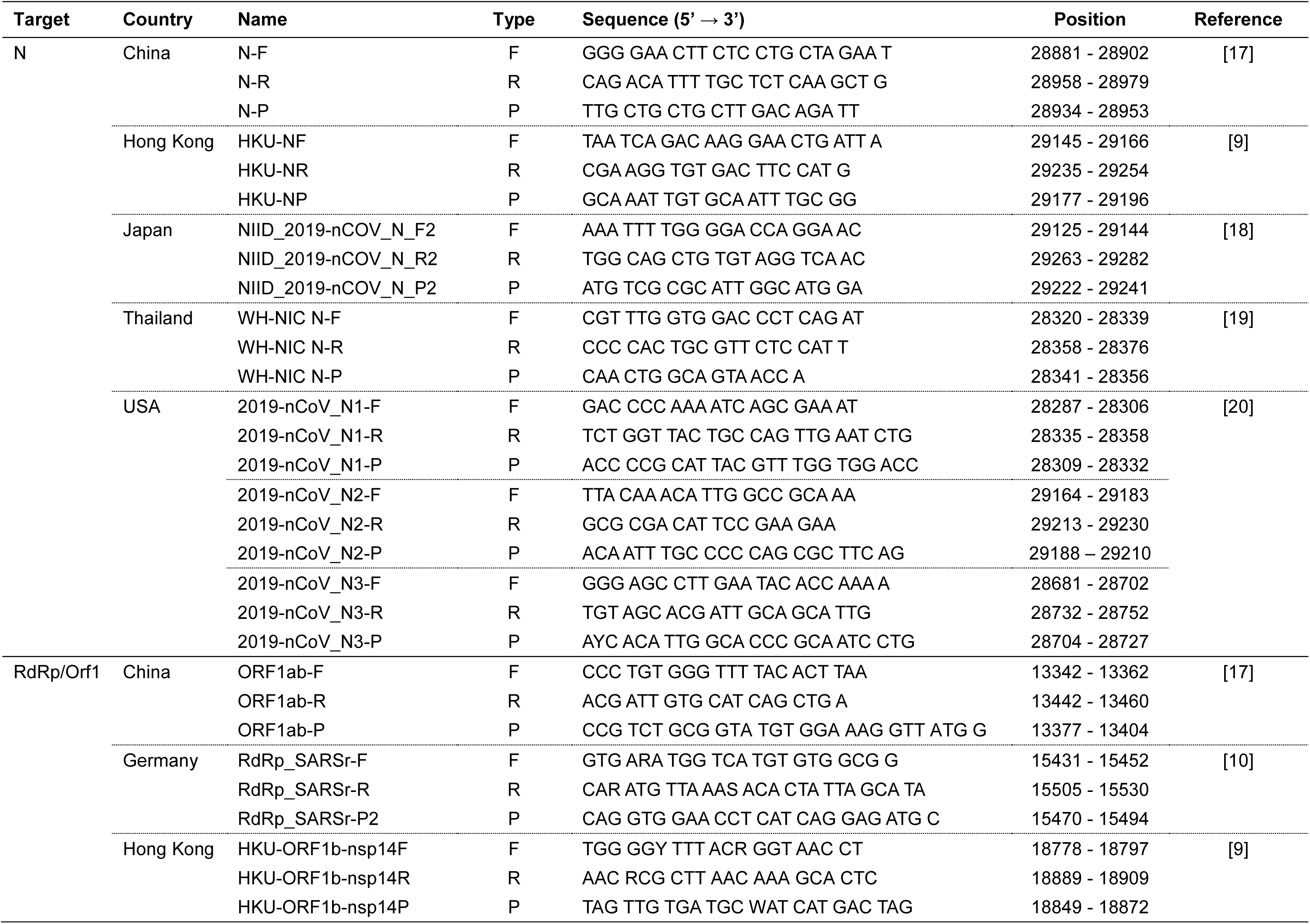
Information of primers and probes analyzed in the study.

**Figure 1.**
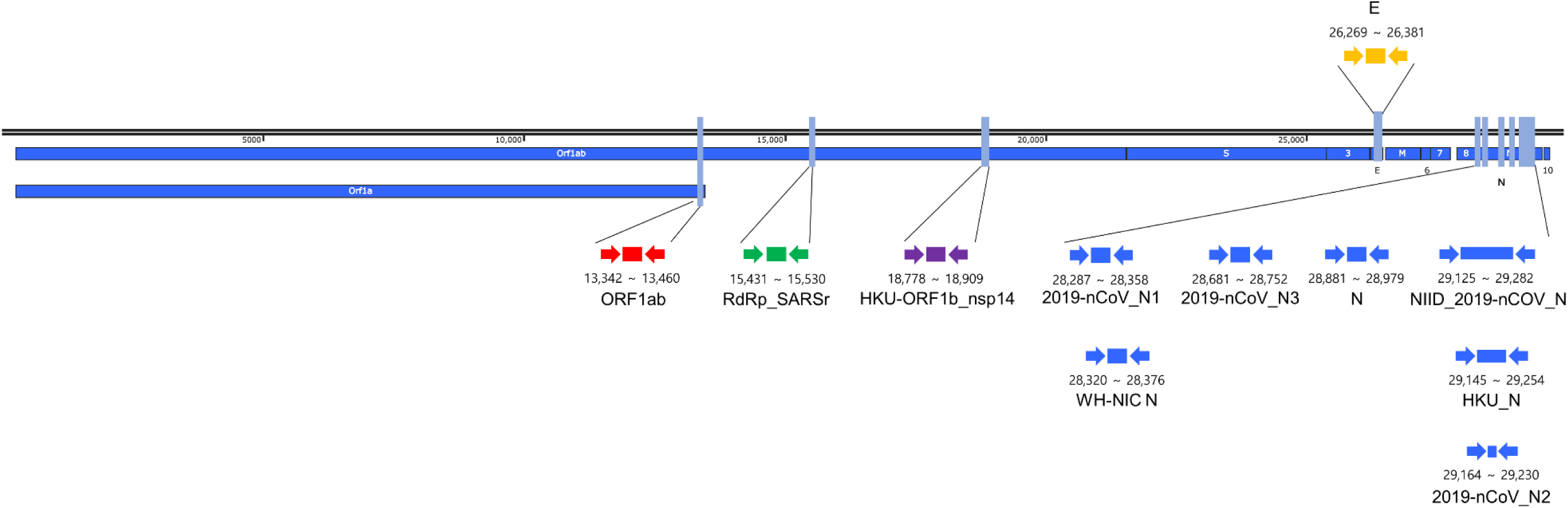
Relative positions of qRT-PCR primer-probe set on the SARS-CoV-2. The target genes and sequences of primers were searched from WHO website (http://www.who.int). The number below amplicons are genome positions according to SARS-CoV-2, GenBank MN908947.3. The sets were published by China CDC (Orf1ab and N), Charité – universitätsmedizin berlin institute of virology in Germany (RdRp_SARSr and E), the University of Hong Kong (HKU-ORF1b_nsp14 and HKU-N), USA CDC (2019-nCoV_N1, N2, and N3), National Institute of Health in Thailand (WH-NIC N), and National Institute of Infectious Disease in Japan (NIID_2019-nCoV_N). Orf1: open reading frame 1; RdRp: RNA-dependent RNA polymerase gene; Nsp14: non-structural protein 14 gene; S: spike protein gene; E: envelop protein gene, N: nucleocapsid protein gene

### Viral RNA preparation

The infection experiments were performed in a biosafety level-3 (BSL-3) laboratory. African green monkey kidney Vero cells (ATCC CCL-81) were infected with a clinical isolate SARS-CoV-2 (BetaCoV/Korea/KCDC03/2020 provided from Korea CDC). After 72 h, the culture medium containing mature infectious virions (virus medium) was collected and viral RNA was isolated from the culture medium using the QIAamp viral RNA extraction Kit (Qiagen, Hilden, Germany) according to the manufacturer’s instructions.

### Preparation of in vitro transcribed RNA standard

The coding sequence of SARS-CoV-2 Envelope (E) protein, which cloned in pET21a plasmid was PCR amplified with T7 promoter primer (5’ – AATACGACTCACTATAG – 3’, Macrogen Inc., South Korea) and T7 terminator primer (5’ – GCTAGTTATTGCTCAGCGG – 3’, Macrogen) with AccuPower® PCR PreMix (-dye) kit (Bioneer Inc., South Korea). PCR product was then used as *in vitro* transcription template using MEGAscript™ T7 Transcription Kit (Invitrogen Inc., CA, USA). The copy number of *in vitro* transcribed RNA was calculated from RNA concentration measured with Quantus™ Fluorometer (Promega Inc., WI, USA). Standardized amounts of *in vitro* produced RNA were used E primer and qRT-PCR to produce a standard curve.

### Confirmatory qRT-PCR in RdRp and N

Extracted nucleic acid samples were tested for comparative analysis of SARS-CoV-2 by qRT-PCR. The Orf1 and N region of SARS-CoV-2 were used as the target sequences for SARS-CoV-2 specific gene. Briefly, 10 μL of purified viral RNA was amplified in a 20 μL reaction solution containing 1X 1 step RT-PCR mix (WELLS BIO INC., South Korea), and 300 nM of primers and probes for the target detection. The qRT-PCR was performed with a CFX 96 touch real-time PCR detection system (Bio-rad, Hercules, CA, USA). The qRT-PCR conditions applied in this study were programmed as follows: UNG incubation, RT incubation, and enzyme activation were serially performed at 25 °C for 2 minutes, at 55 °C for 10 minutes, at 94 °C for 3 minutes respectively. Thermal cycling was then performed at 94 °C for 15 seconds (denaturation), and at 60 °C for 30 seconds (annealing and amplification) for forty-five cycles.

## Results and Discussion

### Validation of qRT-PCR assay

The Ct value was not produced from negative control, indicating the reaction was done aseptically. The standard curve from E gene primer-probe set also showed the reaction was done accordingly. The R^2^ value of the standard curve was 0.999 and the calculated amplification efficiency was 101.6%. These indicated that the qRT-PCR reaction was done in optimal condition. The viral concentration of supernatant and cell lysate was determined by E gene-based assay (Table 2).

**Table 2.** Comparative analysis of Ct values obtained by employing each primer-probe set.

| Target | Country | Name | Ct value |  |  |  |
| --- | --- | --- | --- | --- | --- | --- |
| | | | $1.5 \times 10^4$<br>copies | $1.5 \times 10^3$<br>copies | $1.5 \times 10^2$<br>copies | $1.5 \times 10^1$<br>copies |
| N | China | N | 24.01 | 26.96 | 30.46 | 34.86 |
|  | Hong Kong | HKU-N | 26.00 | 29.45 | 33.17 | 35.43 |
|  | Japan | NIID_2019-nCoV_N | 23.09 | 26.56 | 29.5 | 33.13 |
|  | Thailand | WH-NIC N | 28.64 | 31.89 | 35.26 | 38.13 |
|  | USA | 2019-nCoV_N1 | 24.25 | 27.50 | 30.57 | 34.71 |
|  |  | 2019-nCoV_N2 | 22.88 | 26.12 | 29.26 | 33.14 |
|  |  | 2019-nCoV_N3 | 22.64 | 26.01 | 29.42 | 33.09 |
| RdRp/Orf1 | China | ORF1ab | 27.33 | 30.33 | 33.61 | 36.85 |
|  | Germany | RdRp_SARSr | 31.89 | 35.14 | 38.57 | -* |
|  | Hong Kong | HKU-ORF1b-nsp14 | 29.04 | 32.03 | 35.33 | 38.97 |

\* The assay with RdRp\_SARSr (Germany) set showed a positive signal (43.00) from the single reaction of triplicate.

### RdRp/Orf1 Assays

The Ct value of RdRp_SARSr (Germany), HKU-ORF1b-nsp14 (Hong Kong), and ORF1ab (China) from low concentration (15 copies/reaction) were 43.00, 38.97, and 36.85, respectively (Table 2). The assay with RdRp_SARSr (Germany) set showed a positive signal from the single reaction of triplicate in the concentration of 15 copies/reaction. The assay with HKU-ORF1b-nsp14 (Hong Kong), and ORF1ab (China) sets showed positive signals in the concentration of 1.5 copies/reaction (data not shown). The R^2^ value from RdRp_SARSr (Germany), HKU-ORF1b-nsp14 (Hong Kong), and ORF1ab (China) were 0.983, 0.997 and 0.997, respectively. The calculated amplification efficiency of RdRp_SARSr (Germany), HKU-ORF1b-nsp14 (Hong Kong), and ORF1ab (China) was 101.6%, 96.1%, and 109.8%, respectively. As a result, ORF1ab (China) set may be recommended for the laboratory confirmation of the RdRp/Orf1 gene.

### N Assays

The Ct value of N (China), HKU-N (Hong Kong), NIID_2019-nCOV_N (Japan), WH-NIC N (Thailand), 2019-nCoV_N1, N2, and N3 (USA) from low concentration (15 copies/reaction) were 34.86, 35.43, 33.13, 38.13, 34.71, 33.14, and 33.09, respectively (Table 2). The Ct value of 2019-nCoV_N2, N3 (USA), and NIID_2019-nCOV_N (Japan) sets were similar to each other, and the sets could be regarded as the most sensitive sets. The moderately sensitive assay was based on 2019-nCoV_N1 (USA) and N (China). These sets had higher Ct value than the most sensitive sets, however, the Ct values from low concentration (15 copies/μl) were still within the cut-off value (Ct <37). WH-NIC N (Thailand) set was less sensitive than other sets. The Ct value from low concentration (15 copies/μl) was close to the cut-off value (Ct <38). The R^2^ value from of N (China), HKU-N (Hong Kong), NIID_2019-nCOV_N (Japan), WH-NIC N (Thailand), 2019-nCoV_N1, N2, and N3 (USA) were 0.989, 0.980, 0.987, 0.987, 0.986, 0.952, and 0.991, respectively. The calculated amplification efficiency of N (China), HKU-N (Hong Kong), NIID_2019-nCOV_N (Japan), WH-NIC N (Thailand), 2019-nCoV_N1, N2, and N3 (USA) were 89.4, 105.3, 100.7, 106.2, 95.2, 97.3, and 93.9, respectively. Therefore, 2019-nCoV_N2, N3 (USA), and NIID_2019-nCOV_N (Japan) sets should be beneficial for the laboratory confirmation of SARS-CoV-2 by qRT-PCR assay of N gene.

## Conclusions

Various primer-probe sets were previously reported to detect SARS-CoV-2 by the qRT-PCR assay. The sensitivity of the assay may not enough to confirm suspicious patients in the early stage of SARS-CoV-2 infection. Nevertheless, there have been no comparative results of the current qRT-PCR analysis for the molecular diagnosis of SARS-CoV-2. In the present study, the first comparative analysis of various primer-probe sets targeting RdRp/Orf1 and N region of SARS-CoV-2 was performed by qRT-PCR for the laboratory confirmation. In the case of targeting RdRp/Orf1 region, ORF1ab (China) set might be the most sensitive than other sets. 2019-nCoV_N2, N3 (USA), and NIID_2019-nCOV_N (Japan) sets may be recommended for the sensitive qRT-PCR assay of N region. Therefore, the appropriate combination from ORF1ab (China), 2019-nCoV_N2, N3 (USA), and NIID_2019-nCOV_N (Japan) sets should be selected for the sensitive and reliable laboratory confirmation of SARS-CoV-2.

## Acknowledgments

We appreciated to National Culture Collection for Pathogen of Korea CDC for providing clinical SARS-CoV-2 isolate. This work was supported by National Research Council of Science and Technology grant by the Ministry of Science and ICT (Grant No. CRC-16-01-KRICT).

